# Towards revealing the transient dynamics in plant biomass allocation pattern

**DOI:** 10.1101/2023.05.19.541549

**Authors:** Renfei Chen

## Abstract

1. In addition to allometric biomass partitioning theory, optimal partitioning theory is one of the most important theoretical frameworks in predicting plant biomass allocation patterns. However, focusing on the equilibrium state leads to a mismatch between some empirical observations and estimations from optimal hypotheses.
2. To address this issue, I developed a heuristic approach with a quantitative metric to study the transient patterns of plants allocating photosynthetic products to various combinations among plant organ parts. Moreover, the approach also discovers the mechanisms under which various factors drive the transient patterns.
3. With this approach, I provide a case study and find that the perturbations of the transient patterns of plant leaf and stem biomass periodically decrease and increase in response to plant height, crown diameter, and projected crown area. Predictions with the approach are well demonstrated by empirical data consisting of global forest plants.
4. ***Synthesis***. The approach here complements the limitations of optimal partitioning theory by revealing the variations of plant photosynthetic partitioning in short time scales. Given the central role of plant biomass allocation pattern in both empirical applications and theoretical foundations, there is a large scope for using this approach to investigate the directions in estimations of carbon stock, stabilized yields in agriculture as well as forest management.

## Introduction

An important topic in ecology is how resources absorbed from the environment are allocated to different plant organ parts (Bazzaz and Grace, 1997; Poorter et al., 2015, 2012; Puglielli et al., 2021). The underlying mechanisms have important applications in both basic and applied sciences. For example, estimating carbon stock contributed by plants, especially by plant below-ground (roots) organs which usually is much difficult to measure in practice (Robinson, 2004; Mokany et al., 2006; Siddiq et al., 2021); improving yield in agriculture depending on plant plasticity affected by environmental factors such as soil nutrients and climate change (Weiner, 2004; Tamagno et al., 2018); developing general theoretical frameworks based on the predictions of the fourth dimension of life (West et al., 1999; Enquist and Niklas, 2002). In the end, quite an amount of work has focused on the patterns of photosynthetic resources that are allocated to different plant organs (e.g., plant leaf, stem, and roots) in response to both biotic and abiotic factors (Chen et al., 2019, 2021; Jevon and Lang, 2022). Meanwhile, theories have been developed based on suitable assumptions (Enquist and Niklas, 2002; Peng et al., 2022; Gedroc et al., 1996; Jevon and Lang, 2022). Among which is the optimal allocation theory based on the assumption that plants allocate resources among organs to maximize plant biomass production according to economic analogs (Bazzaz and Grace, 1997; Bloom et al., 1985). In contrast to allometric or isometric allocation theory which believes that environmental factors do not affect the general scaling exponents (Enquist and Niklas, 2002), optimal allocation theory suggests that plants allocate more biomass to the organ parts that suffer the most stress from the environmental factors to maximize growth rate (Bloom et al., 1985; Jevon and Lang, 2022; Chen et al., 2021). The predictions are strongly consistent with many phenomena observed in natural systems. For example, plants in deserts have developed root systems through maximizing water and nutrient uptake to adapt to arid and barren soil environments (Sun et al., 2021; Chen et al., 2019). In contrast, plants living in dark environments allocate more photosynthetic products to plant leaf parts to maximize plant growth through maximizing light resources (Sun et al., 2021; Umaña et al., 2021; McCarthy and Enquist, 2007).

Although optimal allocation theory plays a powerful role in predicting plant biomass allocation patterns, ecologists are increasingly paying attention to its limitations when some empirical data cannot be explained by optimal allocation theory (Lohier et al., 2014; Puglielli et al., 2021). By considering resource limitations that both plant above- and below-ground parts suffer, experiments examining the seedlings from tropical rain forests in China indicate that the allocation strategies between photosynthetic and non-photosynthetic tissues and roots over stems do not always follow the allocation patterns as predicted by the optimal allocation theory (Umaña et al., 2021). Likewise, a precipitation manipulation experiment conducted in semiarid steppe suggests that plants alter their root vertical distributions rather than biomass allocation to adapt to drought stress, and predictions from optimal allocation theory can only be tested when plants survive under extreme drought conditions (Zhang et al., 2019). The contrast conclusions drawn from different empirical observations limit the predictability of optimal hypotheses in plant biomass allocation patterns, which needs to develop new theoretical frameworks. To solve the problem, Lohier et al. (2014) suggest that the naturally dynamic environment factors and potential disturbances as well as initial developmental constraints lead to variable and unequal limitations to shoot and root activities over time, which conflict with the assumptions that optimal allocation theory based on (Lohier et al., 2014). Therefore, the transient dynamics of plant growth can effectively explain the conflicts between theoretical predictions from optimal allocation hypotheses and experimental observations (Lohier et al., 2014).

The significance of studying the transient ecological dynamics relies on the various time scales observed in natural systems (Morozov et al., 2020; Hastings et al., 2018; Hastings, 2010). Relative to long-term scales such as redwood trees or long-lived reptiles which can survive with generation times about one hundred years or even longer, some ecological processes turn out to be very short such as single-celled organisms reproducing within hours and univoltine insects reproducing yearly(Hastings, 2010). Growing studies have demonstrated that population behaviour in asymptotic long time scales is much different from that under short time scales (Hastings and Higgins, 1994; Morozov et al., 2020; Francis et al., 2021), which results in the definitions of “transient dynamics” (Hastings, 2010). In such a case, theories based on long-term asymptotic states cannot match empirical observations that only happened very shortly (Hastings, 2004). Insisting on the analyses only from long time scales can cause not only ecological theories with large limitations in predictions but also destructive damages in empirical applications as sudden changes and regime shifts in transient dynamics can cause population collapse and species extinction (Morozov et al., 2020; Rocha et al., 2018). In the end, transient dynamics have been investigated in many specific areas in ecology (Francis et al., 2021; Shriver et al., 2019). For example, oscillations in transient dynamics can result in species coexistence in either coupled predator-prey systems (Hastings, 2001) or systems consisting of many species that compete for the limiting resources (Huisman and Weissing, 1999); studies about the intertidal salt marshes suggest that transient patterns of fairy circle can not only infer corresponding ecological mechanisms but also reveal the resilience of self-organized systems (Zhao et al., 2021); last but not least, achievements can also be seen in the transient response of fisheries management with the implementation of marine reserves (Chen, 2020; Chen et al., 2022; White et al., 2013).

However, little research can be seen on the transient dynamics of plant biomass allocation patterns. The reason why the research gap exists can be explained by the fact that theories about plant biomass allocation patterns focus on the different organ parts of an individual plant (Enquist and Niklas, 2002; Poorter et al., 2012), while theories about transient dynamics generally aim at populations or communities in most cases (Hastings, 2001; Hastings et al., 2018; Shriver et al., 2019). However, plant components may have similar dynamics as that at the population or community level. For example, the trade-off exists when plants allocate photosynthetic products to leaves, stems, and roots, and thus each organ cannot achieve too much or too few resources to maintain equivalent function (such as absorbing nutrients and photosynthesis) in plant growth. That is the “coexistence among organs”, which is much similar to the case when two or more species struggle to coexist under competition for the limited food resources (Hastings et al., 2018; Huisman and Weissing, 1999). Therefore, it is possible that, during plant ontogeny, the transient dynamics of different plant organs may exhibit contrasting outcomes relative to asymptotic long-term behaviours just like the implications from the transient theories about the dynamics of a population or community. Indeed, Lohier et al. (2014) demonstrated that this ideology with transient analyses is feasible. However, until recently, it is still lacking in developing a quantitative metric as well as a systematic theoretical framework that predicts the transient patterns of plant biomass allocation.

To better understand the extent to which photosynthetic resources are allocated to different plant organs in short time scales, I developed a heuristic framework for quantifying the transient patterns of plant biomass allocation. Specific steps are provided to perform the transient analyses, especially for calculating the transient metric *r* which quantitatively measures the perturbations of a system. With the general procedure, one could investigate the transient dynamics of two interests of plant organs (such as two organ parts among leaf, stem, and root; photosynthetic vs. non-photosynthetic parts; above-vs. below-ground biomass; reproductive vs. vegetative organs) in response to the target influencing factors (biotic factors such as plant ontogeny, plant height and abiotic factors such as precipitation and temperature). Last, the approach is applied into a case study to predict how plant height, as well as crown diameter and projected crown area, regulates the transient patterns of plant leaf and stem biomass, and the theoretical predictions from both analytical and simulation analyses are tested by global empirical forest data.

### Analytical analyses

Here, I use a case study to specifically exhibit how to perform the transient analyses of plant biomass allocation patterns. Firstly, I investigate the transient perturbations of plant leaf and stem biomass in response to plant height. Then, using similar steps, I study the effect of crown diameter and projected crown area as well. As suggested by previous viewpoints (Erickson, 1976), I use trigonometric functions to define the growth rate of plant leaf (*M_L_*) and stem (*M_S_*) biomass.

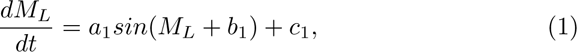

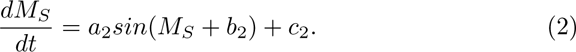

where *a_i_*, *b_i_*, *c_i_* (*i* = 1, 2) are constant parameters. Allometric theories have shown statistically significant scaling relationships between leaf biomass vs. plant height and stem biomass vs. plant height with relatively constant scaling exponents (Niklas, 2003). To be general, I assume that these scaling exponents are unknown parameters in the model, whose specific values are subsequently estimated by the empirical forest data. Therefore, we have

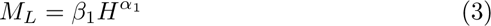

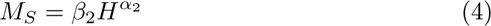

where *α_i_* (*i* = 1, 2) is scaling exponents, and *β_i_* is constants. *H* denotes plant height. Based on Eqs. 7-10, the Jacobian matrix of the system is expressed as a function of plant height

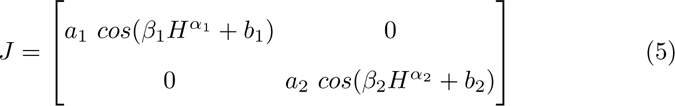

Using Eq. 6, the relationship between transient metric *r* and plant height *H* is

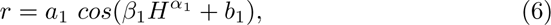

and thus the derivative of g-reactivity *r* in terms of plant height *H* is

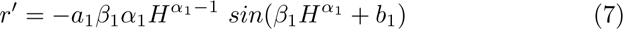

The analytical analyses show that transient metric *r* periodically increases and decreases in response to increasing plant height. In an ecological context, it suggests that, on one hand, the deterministic dynamics of plant leaf and stem biomass during the whole plant ontogeny are much more difficult to predict due to the perturbations caused by plant height rather than implications suggested by optimal allocation theory. On the other hand, it predicts that there is a couple of optimal plant height which can maximally stabilize the dynamics of plant leaf and stem biomass. This has important implications such as applications in achieving stabilized net primary production of trees in forests and yields in agriculture. Similarly, it is feasible to analytically investigate the effect of crown diameter as well as the projected crown area on the transient perturbations of plant leaf and stem biomass, which also shows periodical variations.

### Numerical and empirical results

Consistent with the analytical results, numerical analyses show that transient metric *r* varies periodically in response to the increase in plant height (Fig. 1). Therefore, plant height can either promote or remove the perturbations of plant biomass allocated to both leaf and stem during the transient time scales. Note that the numerical results are based on the assumptions of the allometric relationships between leaf biomass vs. plant height, and between stem biomass vs. plant height. Our regression analyses of the empirical data verified these assumptions. The scaling exponent of plant leaf biomass (*M_L_*) vs. plant height (*H*) is 2, while it is 8/3 for the allometric relationship between plant stem biomass (*M_S_*) and plant height (Fig. 2). These scaling relationships are robust for all the situations when the filtrated global data are either pooled together (Fig. 2A and B) or separated into deciduous (Fig. 2C, D) and evergreen (Fig. 2I, J) plants as well as angiosperm (Fig. 2E, F) and gymnosperm (Fig. 2G, H) plants.

**Fig. 1.**
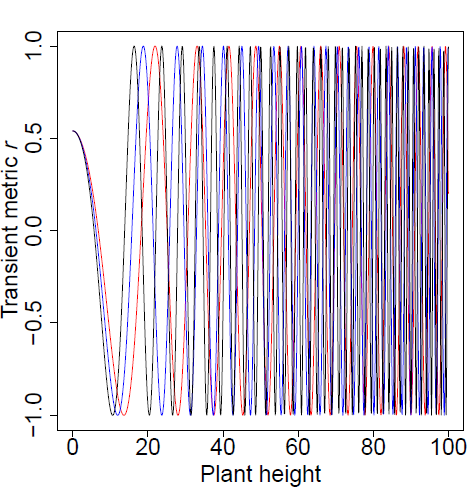
Theoretical predictions of transient metric *r* in response to plant height. Parameters are *a*_1_ = 1, *b*_1_ = 1, *β*_1_ = *e^−^*^4.2^, *α*_1_ =1.9 (red), 2 (blue), 2.1 (black).

**Fig. 2.**
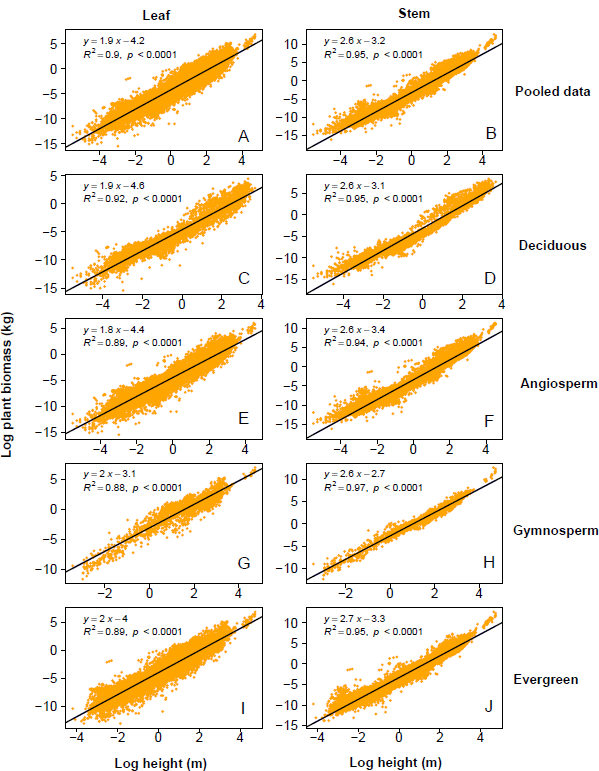
Scaling relationships between plant biomass and plant height with pooled data as well as different plant functional types. For all filtrated, deciduous, angiosperm, gymnosperm, as well as evergreen forest plants, the linear relationships are significant and robust between log-transformed leaf biomass vs. log-transformed plant height (A, C, E, G, I) and between log-transformed stem biomass vs. log-transformed plant height (B, D, F, H, J).

The regression analyses of the empirical data show that the coefficients of variation of leaf biomass, and stem biomass as well as they both increase and decrease periodically in response to different plant height gradients (Fig. 3). Moreover, the periodical variation trends are general among different plant functional types (Fig. 3 *A*_2_ *− A*_5_, *B*_2_ *− B*_5_, *C*_2_ *− C*_5_). Similarly, there are statistically significant allometric relationships between plant biomass and crown diameter as well as the projected crown area (Fig. 4), which regulate the instability of the system consisting of plant leaf and stem (Fig. 5). Therefore, theoretical predictions from both analytical and numerical results are tested by the empirical observations from the global forest data set (Fig. 2-5).

**Fig. 3.**
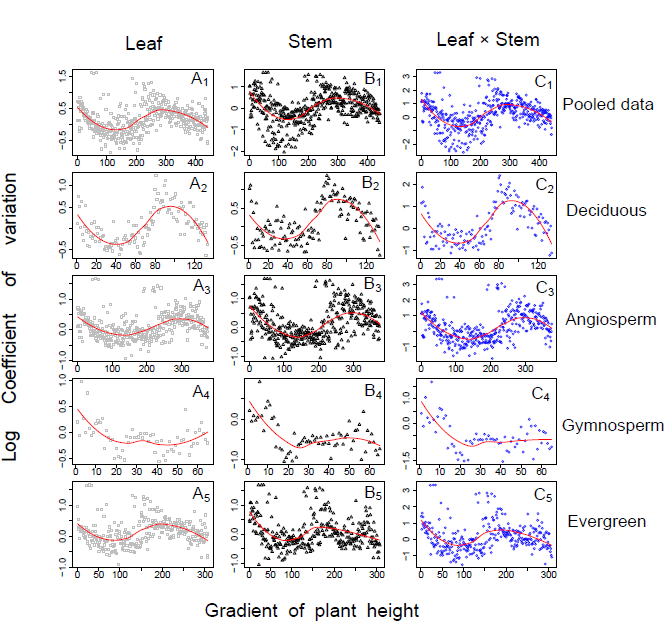
Empirical observations of the variations of plant biomass in response to the gradient of plant height. Under different plant height gradient, the coefficients of variation of leaf biomass (*A*_1_ - *A*_5_), stem biomass (*B*_1_ - *B*_5_) as well both (*C*_1_ - *C*_5_) oscillates for all filtrated, deciduous, angiosperm, gymnosperm, as well as evergreen forest plants. The red lines denote the variation trends based on the Local Polynomial Regression analyses.

**Fig. 4.**
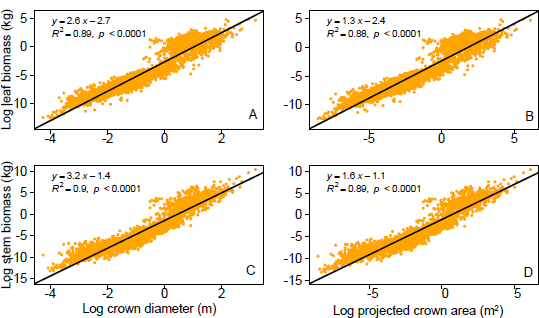
Empirical observations of scaling relationships between plant biomass of different organs and plant traits. (A) leaf biomass vs. crown diameter, (B) leaf biomass vs. projected crown area, (C) stem biomass vs. crown diameter, and (D) stem biomass vs. projected crown area.

**Fig. 5.**
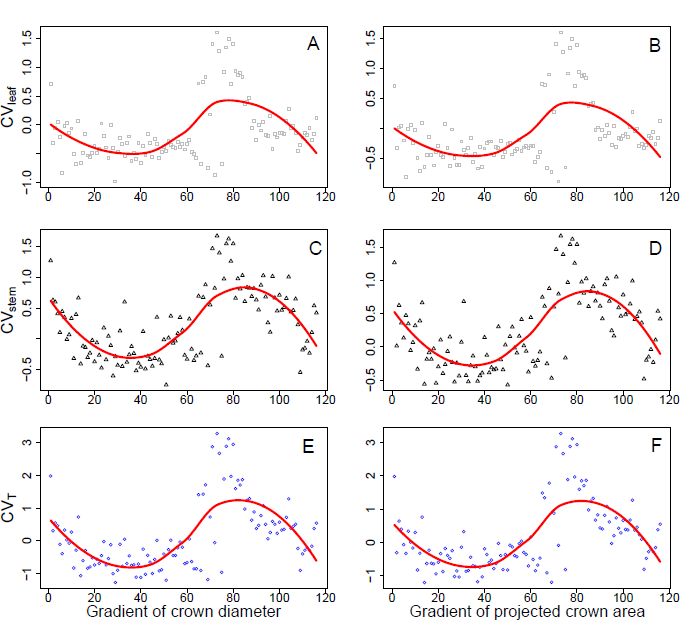
Empirical observations of coefficient of variations in response to crown size. Variations of plant leaf biomass (A, B), stem biomass (C, D), and both (E, F) in response to the gradient of crown diameter (A, C, E) and projected crown area (B, D, F). The red lines denote the variation trends based on the Local Polynomial Regression analyses.

## Discussion

By integrating transient metric and allometric theory, my research provides a quantitative approach to investigating the extent to which plant biomass allocation patterns can oscillate during plant ontogeny in transient time scales. Moreover, it helps us understand how the perturbations of different plant organ parts are regulated by various biotic and environmental factors. Transient phenomena are universal in ecological systems (Hastings et al., 2018), which can also be observed in plant biomass allocation patterns as suggested by my research. Therefore, rather than focusing on the equilibrium state conditions that the well-known optimal allocation theory is based on, the approach here can be used to discover the transient pattern of plant biomass allocation from a quantitative perspective. This can provide theoretical foundations for some important and contrasting phenomena that cannot be explained by the classic optimal allocation theory, and provide practical applications in stabilizing yields of various ecosystems. The approach is based on general theoretical frameworks, and thus can be applied to most of the specific scenarios in the biomass allocation pattern between two plant organ parts.

To clearly show how to perform the analysis procedures of the approach, I provide case studies about the transient pattern of plant leaf and stem biomass in response to plant height and crown size. I find that all of these influencing factors (plant height and crown size) can periodically regulate the transient perturbations of both leaf and stem biomass. In other words, the instability of plant biomass that is allocated to different organ parts periodically increases or decreases. These findings have important theoretical implications as the transient biomass allocation patterns complement the predictions of optimal allocation theory from long-term scales. On a long-term scale, plants suffering from environmental stress (e.g. lack of light resources) tend to allocate more biomass to above-ground growth according to optimal allocation theory (Gedroc et al., 1996; Sun et al., 2021; Umaña et al., 2021; McCarthy and Enquist, 2007), and this phenomenon may be observed in equilibrium state after the implementation of the dark stress for a long time. However, in transient time scales, plant allocation patterns change “on the way” which may not be the same as the pattern shown at the long-term equilibrium state. Therefore, these findings can be used to explain why some empirical observations cannot match predictions from optimal allocation theory.

Although a large number of studies on plant biomass allocation patterns still focus on testing either isometric theory (Yang et al., 2009; Eziz et al., 2017; Liu et al., 2021) or optimal allocation theory (Xie et al., 2016; Freschet et al., 2018; Gao et al., 2021) or both, some studies or implications pay attention to the transient dynamics of plant biomass allocation pattern. As suggested, plant growth, as well as its allocation pattern, can change across many potential relationships (Robinson, 2020). Under ontogenetic developmental constraints and nutrient limitations, results on root shoot allometries suggest that insight into plant transient allocation patterns enables empirical observations to match the theoretical predictions from optimal allocation theory (Lohier et al., 2014). Here, my approach extends the analyses from root shoot relationships to more general relationships among different plant organs (including but not limited to leaf, stem, and root). Moreover, the transient metric that my approach provided entails quantitative estimations of the extent to which plant organs can oscillate in response to either environmental or intrinsic factors.

My approach also has important empirical applications such as in forest and agricultural management. As plant allocation patterns to leaf biomass are strongly related to assimilation capacities, and thus net primary production (Chen et al., 2021), my approach can be used to serve forest management by stabilizing net primary production. According to my analyses, plant traits such as plant height and crown size can produce periodical effects on the perturbations of biomass allocated to the leaf (Figs. 3 and 5). An important implication is that observations on easily measured plant height and crown size can predict whether and when we can achieve stabilized net primary production for a certain forest. Likewise, in agronomy, farmers not only pay attention to high yields but also focus on yield stability which is a concept that has no complete agreement on its definition, and an intuitive and widely accepted definition is the inverse of the variation in yield (Boussakouran et al., 2021; Tollenaar and Lee, 2002; Weiner et al., 2021). To improve yield stability, large amounts of efforts have been dedicated to maintaining the survival and growth of crops while ignoring the allocation pattern to plant reproduction parts which are the yield for most crops (Weiner et al., 2021). My approach here can also be used to predict the transient allocation pattern between vegetative and reproductive biomass as well as how this pattern is controlled by biotic and abiotic factors. For example, by replacing leaf and stem biomass in Eqs. 7 and 8 with vegetative and reproductive biomass, a simple analytical analysis suggests that the stability of biomass allocating to plant reproductive parts and thus crop yields periodically vary in response to plant height. Therefore, we could choose the optimal plant height through genetic selection or environmental control to maximize yield stability. Moreover, plant height positively correlated with plant stem biomass (Fig. 2; (Niklas, 2003)), which suggests that a higher plant allocates more biomass to plant stem organ parts and thus lesser to other organs including reproductive parts. In such a case, plant height may affect maximum crop yield. Indeed, studies have suggested that plant features and traits can greatly regulate crop yield (Mathan et al., 2016). In summary, plant height can simultaneously regulate maximum crop yield as well as yield stability, and my approach is useful to achieve the two goals by predicting the optimal plant height for yield stability constrained in the spectrum of plant heights that have been suggested to maximize crop yield.

An important assumption I used here is the allometric scaling relationships between the interested plant organ parts and their influencing factors. The analytical analyses suggest that the periodical relationship between transient metric and drivers are robust and general regardless of the specific value of the scaling exponents. Therefore, when using the empirical data to test the theoretical predictions, it is more important to pay attention to the goodness of fit rather than the scaling exponents although scaling exponents play an important role in allometric theories as well as in the metabolic theory of ecology (West et al., 1997; Enquist and Niklas, 2001). Because the allometric relationships are universal among different plant organ parts (West et al., 1999; Enquist and Niklas, 2002), it is general enough for us to use the approach to analyze the transient pattern of plant biomass allocation. Future work would be necessary to use this heuristic approach to study more specific biomass allocation patterns not only in cases to test the validity of the approach through empirical observations when variables are easy to achieve from experiments but also in cases to provide theoretical predictions when empirical data are not sufficient.

## Materials and methods

Generally, the procedure to develop the transient pattern analyses needs only four simple steps: i) configuration of the allocation dynamics between plant organs that one wants to study; ii) estimations of the allometric relationships between interested plant organ biomass and influencing factors; iii) analytically calculating transient metrics that denote the perturbations of the system one studied and analyzing its variations in response to influencing factors; iv) concluding and testing the predictions with empirical observations (see details below).

### Dynamic models

First, I use ordinary differential equations (ODE) to develop heuristic models describing the growth rate of two plant interests *m*_1_ and *m*_2_ during plant ontogeny *t*

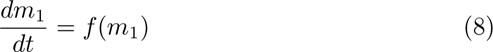

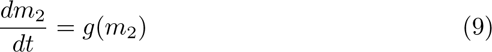

where the general function of *f*() and *g*() could be any specific form such as trigonometric functions. Plant interests *m*_1_ and *m*_2_ are the two contrast parts of plant organs that one wants to study. For example, they could be biomass, area, diameter of plant leaves, stems, and roots as well as the ratios among these plant organs.

### Allometric relationships

Second, using background knowledge to predict how the two plant interests *m*_1_ and *m*_2_ are regulated by influencing factors *I*. For example, the scaling allometric relationships between plant individual biomass and population densities (Enquist and Niklas, 2001; Deng et al., 2012). Thus, we have

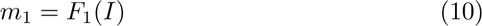

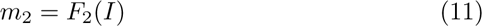

### Transient metric analyses

Third, to analyze how the allocations of different plant organs vary in response to influencing factors in transient time scales, we need to calculate transient metric *r*, which is called reactivity in areas that study population dynamics (Neubert and Caswell, 1997; Mari et al., 2017). Although the specific calculations of reactivity keep on changing with the occurrence of newly developed methods (Mari et al., 2017), its ecological meanings are constant: a high value of transient metric *r* suggests high perturbations of the system. To calculate *r*, we need to calculate the Jacobian matrix of the system (i.e., Eqs. 1 and 2)

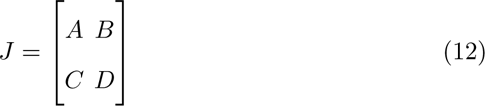

where *A*, *B*, *C*, and *D* are expressions that calculated based on the functions of *f* (*m*_1_) and *g*(*m*_2_) as well as the relationships between *m*_1_ vs. *I* and *m*_2_ vs. *I*. Studies that focus on the transient patterns of population dynamics suggests that it is necessary to firstly define an exemplificative output matrix *c* = [*θ,* 0] to insight into transient metrics (Mari et al., 2017). Consistently, the g-reactivity (a revised version of reactivity) *r* representing the maximum initial amplification rate of perturbations of a system are

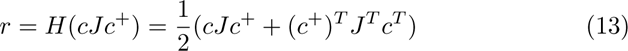

where *H*() is the dominant eigenvalue of the Hermitian matrix, and *c*^+^ = *c^T^* (*cc^T^*)*^−^*^1^ is the right pseudo-inverse of matrix *c*. Substitute equations 5 into equation 6, we could achieve the deterministic relationship between transient metric *r* and parameter *I* denoting influencing factors. Then, the derivatives of *r* with respect to *I* are calculated to predict how influencing factor *I* regulate the perturbations of the system in transient time scales.

### Conclusions and test

With the analytical results, we need to judge the sign of the value of derivatives to draw a conclusion. Generally, the negative value of derivatives of *r* suggests that the perturbations of the system decrease and approximate a stable state with the increasing value of influencing factor *I*. On the contrary, when there is a positive value of derivatives of *r*, the instability of the system increases in response to strengthened influencing factor *I*. Last, numerical and empirical observations of the coefficients of variation (CV) of the interested plant organs are necessary to test the validations of the analytical predictions.

### Numerical approach

To observe how the perturbations of the system vary in response to plant height, I perform numerical analyses revealing photosynthetic allocation to plant leaf and stem biomass. The variations of transient metric *r* with the increasing of plant height are simulated through Eq. 12 derived from analytical solutions. Plant height ranges from zero to one hundred which covers almost all the plants discovered on earth. The numerical analyses are performed in R. 4. 0. 2.

### Empirical test

I tested both the analytical and numerical analyses described above to determine the transient dynamics of plant biomass allocated to leaf and stem with a global database of biomass and allometry for woody plants from published works (Falster et al., 2015). The database consists of plant measurements from individual levels rather than stand averages, and plant heights range from 0.01 to 100 m. Moreover, rather than indirectly predicted from allometric equations, all data about plant biomass were estimated directly. To observe how plant height drives plant biomass allocation patterns in transient time scales, I filtrate the database with the criteria that no absence occurs in the variables of plant leaf biomass, plant stem biomass, and plant height. Under these criteria, I achieve a new database including different plant functional types of evergreen vs. deciduous plants, angiosperm vs. gymnosperm. Using the same criteria, I filtrate the database to study the effect of diameter or width of the crown, and projected crown area as seen from above on the transient allocation to plant leaf and stem, respectively.

With the filtrated database, I conduct statistical regression analyses to predict the allometric relationships between log-transformed plant leaf biomass vs. log-transformed plant height, and log-transformed plant stem biomass vs. log-transformed plant height. The regression analyses are performed with the linear model in the function trendline () in R package basicTrendline. To observe the effect of plant height on the variations of both plant leaf and stem biomass, I perform Local Polynomial Regression with the following steps. First, I rank the data from the lowest to the highest plant height, and then the whole data are sorted into different groups based on plant height gradients. Within each group, I have the same sample size of 30 which has important implications in biostatistics. I use the 30 data in each group to calculate the coefficient of the variations of both plant leaf (*CV_leaf_*) and stem (*CV_stem_*) biomass, respectively. As the variations of both leaf and stem biomass contribute to the total perturbations of the system, I calculate the total variations (*CV_T_*) of the system with the product of the coefficient of the variations of both plant leaf (*CV_leaf_*) and stem (*CV_stem_*) biomass. Therefore, within each group, I achieve three indexes (*CV_T_*, *CV_leaf_*, and *CV_stem_*) denoting the perturbations of the transient dynamics of the system consisting of plant leaf and stem. Last, I marked each group from number 1 to the maximum corresponding to plant height from the lowest to the highest gradients. Scatter plots are drawn to describe the relationships between the coefficient of variations (*CV_T_*, *CV_leaf_*, and *CV_stem_*) vs. group number, and Local Polynomial Regression analyses are performed to estimate the variation trend with the loess() function in R. 4. 0. 2.

## Declarations

- Funding: This work was supported by National Natural Science Foundation of China (grant 32101235) and the Basic Research Program of Shanxi Province (grant 20210302124141) to R.F.C.
- Competing interests: There are no competing interests
- Ethics approval: Not applicable
- Consent to participate: Yes
- Consent for publication: Yes
- Availability of data and materials: the empirical data can be achieved from the references cited. Falster D S, Duursma R A, Ishihara M I, et al. BAAD: A biomass and allometry database for woody plants[R]. Ecological Society of America, 2015.
- Code availability: see appendix for all the code that performed the analyses

